# R2C2+UMI: Combining concatemeric consensus sequencing with unique molecular identifiers enables ultra-accurate sequencing of amplicons on Oxford Nanopore Technologies sequencers

**DOI:** 10.1101/2023.08.19.553937

**Authors:** Dori Z.Q. Deng, Jack Verhage, Celine Neudorf, Russell Corbett-Detig, Honey Mekonen, Peter J. Castaldi, Christopher Vollmers

## Abstract

The sequencing of PCR amplicons is a core application of high-throughput sequencing technology. Using unique molecular identifiers (UMIs), individual amplified molecules can be sequenced to very high accuracy on an Illumina sequencer. However, Illumina sequencers have limited read length and are therefore restricted to sequencing amplicons shorter than 600bp unless using inefficient synthetic long-read approaches. Native long-read sequencers from Pacific Biosciences and Oxford Nanopore Technologies can, using consensus read approaches, match or exceed Illumina quality while achieving much longer read lengths. Using a circularization-based concatemeric consensus sequencing approach (R2C2) paired with UMIs (R2C2+UMI) we show that we can sequence ∼550nt antibody heavy-chain (IGH) and ∼1500nt 16S amplicons at accuracies up to and exceeding Q50 (<1 error in 100,0000 sequenced bases), which exceeds accuracies of UMI-supported Illumina paired sequencing as well as synthetic long-read approaches.

## Introduction

High-throughput sequencing of PCR amplicons is an increasingly essential application that enables a large range of biological, medical, and diagnostic analyses and investigations. Amplicon sequencing can provide detailed insights into the molecular diversity at targeted genetic elements. For example, IGH amplicon sequencing can be used to determine the activation state of the adaptive immune system^1^, track minimal residual disease^2^, or perform basic research into the human immune system^3^. Similarly, 16S amplicon sequencing can be used to profile the composition of bacteria in a variety of biosamples in clinical diagnostics or basic research^4^. For each, high sequencing accuracy is crucial because each sequenced molecule can represent a unique isolate within the sample.

Amplicon sequencing approaches have evolved substantially in recent years and present a host of opportunities for improvement. Illumina sequencers provide good accuracy, but the molecule length limitations complicate the sequencing of amplicons longer than 600bp. For example, a full-length 16S amplicon is ∼1500nt in length and therefore requires synthetic long read approaches like LoopSeq. LoopSeq can achieve accuracies around Q42 but has a complex, proprietary library prep and doesn’t generate many full-length molecules^5^.

Long-read sequencing technologies, Oxford Nanopore Technologies (ONT) or Pacific Biosciences (PacBio), can sequence longer amplicons. Methods like PacBio CCS, and ONT-based Nano-ampliseq and INC-seq have explored concatemer-based consensus approaches for 16S sequencing^6–8^ but generally only reach accuracies below Q30 - far below LoopSeq accuracy. ONT reads generated using the new R10 pore chemistry have been used in conjunction with UMIs to sequence 16S or full-length rRNA operon amplicons and achieve an accuracy matching LoopSeq (∼Q42)^9^. For the same full-length rRNA operons, PacBio CCS and UMIs achieved an accuracy of ∼Q52^9^. PacBio CCS and UMIs have also been used by FLAIRR-seq which analyzed full-length IGH amplicons but did not empirically determine its method’s accuracy^10^. In both cases consensus generation was either done by a multi-step adhoc chain of tools or a likely suboptimal approach designed for Illumina reads.

In summary, while there a several powerful approaches currently available for high-accuracy amplicon sequencing, there currently exists no single approach that 1) is mostly agnostic to amplicon length, 2) has an easy-to-implement and robust library preparation protocol 3) easy-to-use and efficient computational analysis pipeline, 4) achieves accuracies up to and exceeding Q50, and 5) is cost-effective on the instrument and per-read level.

Here, we established this approach by combining our ONT-based R2C2 method^11–16^ with UMI-containing Illumina-style amplicons to generate the easy-to-use R2C2+UMI method. We also introduce the BC1 computational tool (https://github.com/christopher-vollmers/BC1) which we developed to generate R2C2+UMI consensus reads. We used the resulting molecular and computational methods to analyze two amplicon types: A 550nt Illumina-style (p5/p7 adapters) IGH amplicon and a ∼1500nt Illumina-style 16S amplicon. We compared the resulting R2C2+UMI data to either a 2x300 MiSeq flow cell (Illumina+UMI for IGH amplicon) or publicly available LoopSeq synthetic long read data (16S amplicon). We show that R2C2+UMI outperforms the accuracy of both, reaching Q52, thereby establishing new accuracy benchmarks for both IGH and 16S amplicon sequencing.

R2C2+UMI in combination with the BC1 tool enables unprecedented accuracy in interpreting variation captured by amplicon sequencing techniques.

## Results

To achieve the highest possible accuracy for sequencing amplicons, we combined both molecular and computational methods.

First, we used an established protocol^17^ to generate Illumina-style amplicons with each original molecule labeled by an unique molecular identifier (UMI). To sequence these Illumina-style amplicons on ONT long-read sequencers, we processed the libraries into high molecular weight DNA using the R2C2 method. R2C2 circularizes library molecules using Gibson assembly. It then uses rolling circle amplification (RCA) to generate long, linear concatemers containing multiple tandem repeats of the original library molecule.

Second, after sequencing this concatemeric DNA on ONT sequencers, we used the computational C3POa and BC1 tools to generate consensus sequences for each original library molecule. C3POa parses concatemeric raw reads into subreads and generates accurate R2C2 consensus reads from these subreads. BC1 parses R2C2 consensus reads using a highly flexible syntax for the locating and parsing of UMI sequences, enabling the detection of fixed bases used as spacers or IUPAC wildcard base codes like B,D,H,V, R, or Y which, in addition to Ns, can be used to optimize UMIs for more indel-prone long-reads. BC1 then uses the R2C2 consensus reads and subreads associated with each UMI to generate and polish R2C2+UMI consensus reads for each original molecule: R2C2 consensus reads are aligned to each other by abpoa^18^ to form a preliminary consensus. Using the subreads, this preliminary consensus is then error-corrected using racon^19^ and polished using medaka^20^. Because each R2C2 consensus read can be based on multiple subreads (∼4-20 subreads per R2C2 consensus read in this study) and each UMI can be covered by multiple R2C2 consensus reads (e.g. ∼20 R2C2 consensus reads for IGH amplicons in this study), R2C2+UMI consensus reads can be covered by hundreds of subreads and consequently reach very high accuracies.

### IGH sequencing

We applied the R2C2+UMI approach to ∼550nt IGH libraries. We created these libraries as previously described^1,3,17,21–23^ to capture most of the information contained within antibody heavy chain transcripts. In short, following reverse transcription of human whole blood RNA, dsDNA was generated using a UMI-containing primer pool for Framing Region 1 and another UMI-containing primer pool for the first exon of the isotype-defining constant regions (Supplemental Table S1). This dsDNA is then amplified using indexed Illumina p5/p7 primers. This IGH Illumina amplicon library was first sequenced on an Illumina MiSeq and then processed with R2C2 using a protocol we previously developed for Illumina libraries^11^. We sequenced the resulting R2C2 library on a single PromethION flow cell as part of a pool libraries in which it represented about 30% of pooled DNA. We processed the resulting raw reads using C3POa resulting in 8,397,298 R2C2 consensus reads, 6,263,929 of which showed p5/p7 adapters on their ends. C3POa further demultiplexed these reads producing 1,984,245 reads for the library. Because the original Illumina library was only sequenced to read depths of 383,867 MiSeq 2x300 reads, we subsampled the demultiplexed R2C2 reads to this read depth for further analysis.

We then identified UMIs and combined similar UMIs (<=1 mismatch) into groups and generated UMI-based consensus reads using the presto toolkit^24^ for Illumina data and the BC1 tool for R2C2 data. These separate workflows resulted in 23,785 Illumina+UMI consensus reads and 28,511 R2C2+UMI consensus reads. The read coverage of these UMI-based consensus reads were highly similar between the R2C2 and Illumina technologies (Fig. 2A).

**Fig. 1:**
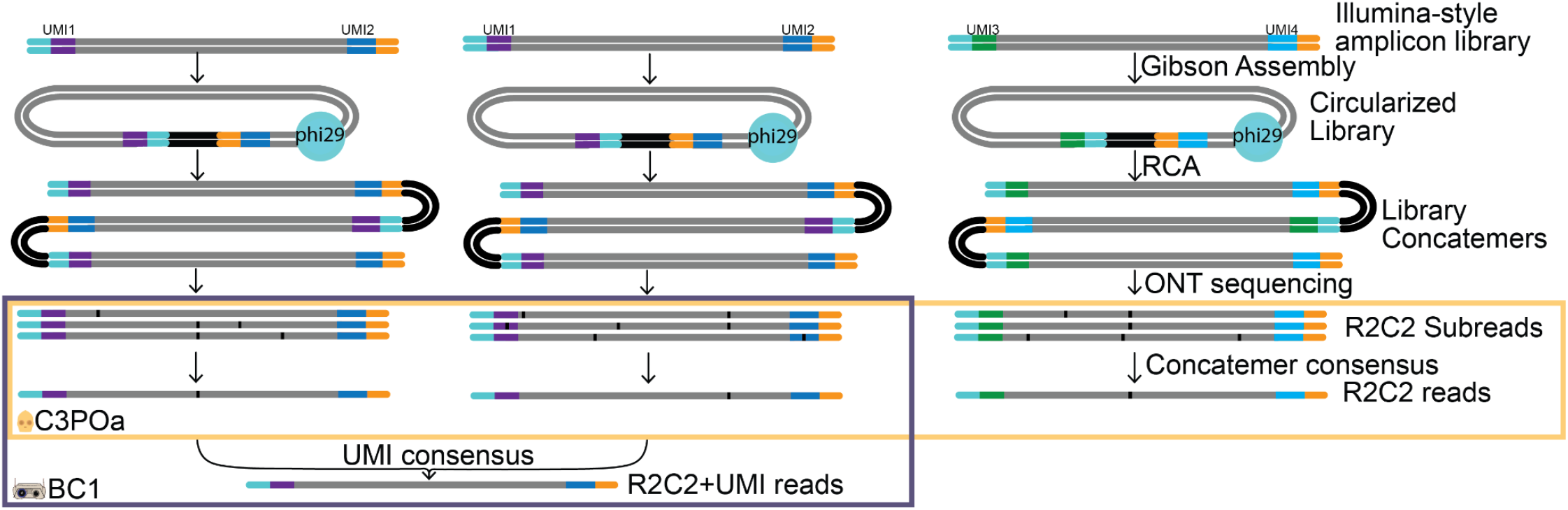
Combining concatemer and UMI based consensus approaches. Illumina-style libraries containing UMIs are converted to R2C2 libraries and sequenced on ONT sequencers. C3POa splits the resulting raw reads into subreads and combines these subreads into high accuracy R2C2 consensus reads. BC1 parses the UMIs contained in these consensus reads, groups reads with identical or highly similar UMIs and uses R2C2 consensus reads and subreads of these groups to generate R2C2+UMI consensus reads of even higher accuracy.

**Fig. 2:**
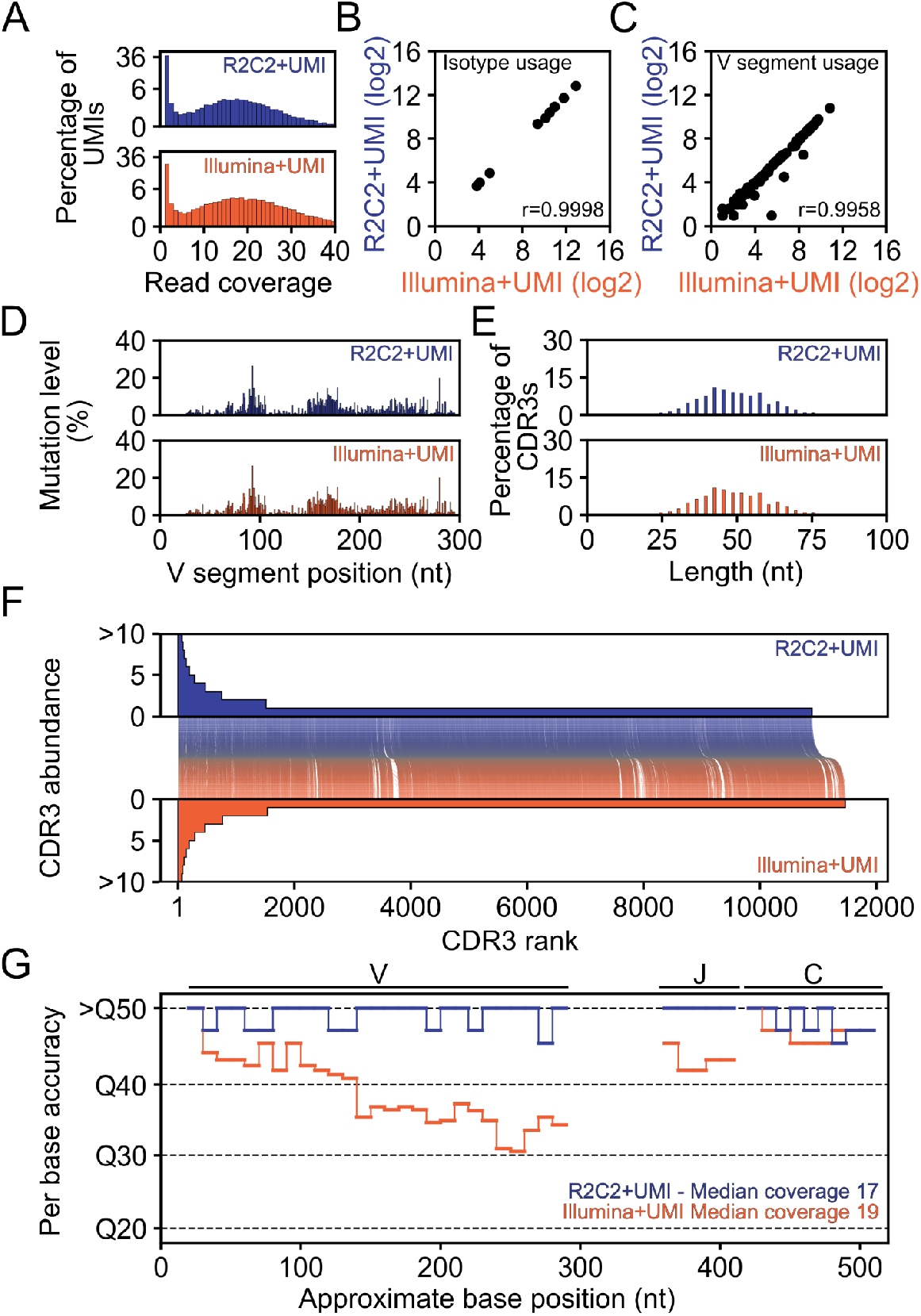
IGH amplicon sequencing. Several characteristics are shown for an UMI-containing IGH amplicon library sequenced either on a Illumina MiSeq (Illumina+UMI) or using R2C2 on a ONT PromethION (R2C2+UMI) A) Read coverage distribution of UMIs (log2 scaled y-axis), B-C) Isotype and V segment usage (log2-converted read numbers) D) Somatic hypermutation location and levels, E) CDR3 length distribution, F) CDR3 abundance and co-occurrence between the two methods, and G) estimated per base accuracy are shown for R2C2+UMI (blue) and Illumina+UMI (orange) approaches

For subsequent analysis, we used IgBlast^25,26^ to process R2C2+UMI and Illumina+UMI reads. IgBlast processing yielded 27,300 R2C2+UMI and 22,620 Illumina+UMI reads that were matched to antibody gene segments. To focus only on highly accurate reads, we restricted further analysis on reads that had four or more Illumina or R2C2 reads covering their UMI - 16,261 Illumina+UMI and 15,743 R2C2+UMI reads. Our analysis covered a comprehensive list of information contained within IGH amplicons. First, we found that Isotype (Fig. 2B) and V segment usage (Fig. 2C) were highly similar between methods. Further, we found that the somatic hypermutation locations and levels within those V segments (Fig. 2D) and the length distribution of CDR3s created by somatic recombination were highly similar between methods (Fig. 2E).

Additionally, the Illumina+UMI and R2C2+UMI methods were used to sequence aliquots of the same IGH library containing copies of the same original molecules which, based on the read coverage distribution of UMIs (Fig. 2A), we infer were sequenced to exhaustion by both methods.

First, this implied that the Illumina+UMI and R2C2+UMI methods should identify identical CDR3 sequences. Indeed, the methods detected 9864 shared CDR3s, however R2C2+UMI detected 1036 unique CDR3s and Illumina+UMI detected 1604 unique UMIs. CDR3s unique to any method suggest sequencing errors in either method changed an existing CDR3 sequence to an artifactual CDR3 sequence.

Second, this implied that the Illumina+UMI and R2C2+UMI methods should identify identical UMIs. This allowed us to estimate Illumina+UMI and R2C2+UMI error rates which otherwise would have been impossible, because IGH genes can be modified by somatic hypermutation and therefore lack an accurate reference. By comparing the Illumina+UMI and R2C2+UMI sequences for the same molecule - as identified by the same UMI - as well as an unmutated reference, we showed that R2C2+UMI consensus reads had an average per-base accuracies of Q51.7 (<1 error: 100,000 sequenced bases) compared to Q37.7 for Illumina+UMI. The comparably low Illumina+UMI average per-base accuracy was surprising so we determined per-base accuracy of both methods throughout the amplicon. Illumina+UMI had comparable accuracy to R2C2+UMI at the amplicon ends, however R2C2+UMI maintained this accuracy throughout the entire amplicon, while Illumina accuracy dips to Q30 in the center of the amplicon. This is likely a consequence of Illumina MiSeq 2x300 paired reads featuring declining accuracies towards their ends. This declining raw accuracy seems to be severe enough that even combining 4 or more Illumina reads cannot create a highly accurate consensus read.

Overall this analysis indicated that the R2C2 preparation did not distort library composition. Further it suggested that R2C2, with the same number of reads, generated equivalent metrics from IGH libraries as Illumina MiSeq 2x300 reads. Importantly, R2C2+UMI consensus reads were 100-times more accurate than Illumina+UMI consensus reads at the same coverage in the center of ∼550nt amplicons.

### 16S sequencing

Next we evaluated the performance of R2C2+UMI for 16S amplicons. 16S amplicons are too long to be sequenced with Illumina paired-end read approaches, both because of read as well as cluster generation length limitations. Because of this, we compared R2C2+UMI reads covering full-length ∼1500nt 16S amplicons generated from the ZymoBIOMICS Microbial Community DNA Standard (Zymo Research D6305) to uncorrected ONT, PacBio, and LoopSeq synthetic long read data generated from the same standard^5,8^.

To this end, we generated 16S dsDNA using previously published primers^27^ modified to contain UMIs (Supplemental Table S1) and then amplified this dsDNA using indexed p5/p7 Illumina adapters. To test whether BC1 can accommodate a wide range of UMI designs we specifically designed these UMIs to not feature homopolymers longer than 3 nucleotides by using a BDHV nucleotide pattern. This UMI design is likely incompatible with Illumina sequencing which requires nucleotide diversity at each position but will make it possible to identify and exclude UMI-containing insertion and deletion errors which are more common in ONT and PacBio data.

We sequenced these amplicons for about 18 hours on a PromethION flow cell resulting in 2,545,474 total R2C2 reads - <25% of this flow cell ultimate total output of ∼15 million reads. 2,319,680 of these R2C2 reads contained p5/p7 adapters and the expected combination of sample indexes. BC1 parsed the custom designed UMIs and consolidated these R2C2 reads into 444,217 reads with unique UMIs - R2C2+UMI reads. 84,154 of these R2C2+UMI reads were covered by more than one R2C2 read. We then used a sliding window approach as previously described^5^ to identify and remove chimeric reads and filtered the remaining R2C2+UMI reads for read length between 1400-1600nt resulting in 77,800 multi-read, non chimeric, and length-filtered R2C2+UMI reads.

36,277 of these filtered R2C2+UMI reads had a coverage of >=12 which we subsequently compared to 17,862 LoopSeq reads. We also included ONT 1D reads derived from R2C2 subreads and PacBio reads^8^ generated from the same ZYMOBIOMICS standard to provide context to this comparison.

To compare read accuracy, we aligned reads to 16S gene reference sequences for the ZYMOBIOMICS standard using minimap2 and then determined errors for each read.

First, we counted the total number of errors produced by the four technologies. Across 10,000 randomly sampled ∼1,500nt reads, R2C2+UMI contained 509 errors, LoopSeq contained 1,570 errors, PacBio CCS reads contained 12,470 errors, and ONT 1D reads contained 90,110 errors.

Second, we calculated the percentage of error-free reads produced by the four technologies. 94.95% of R2C2+UMI reads were free of errors, followed by LoopSeq (92.54%), PacBio (40.44%), and ONT 1D reads (4.72%) (Fig. 3A). The overall number of errors per 10,000 reads as well as the percentage of error-free reads meant that overall accuracies were Q44.8 for R2C2+UMI, Q42.6 for LoopSeq, Q32 for PacBio, and Q24 for ONT 1D reads.

**Fig. 3:**
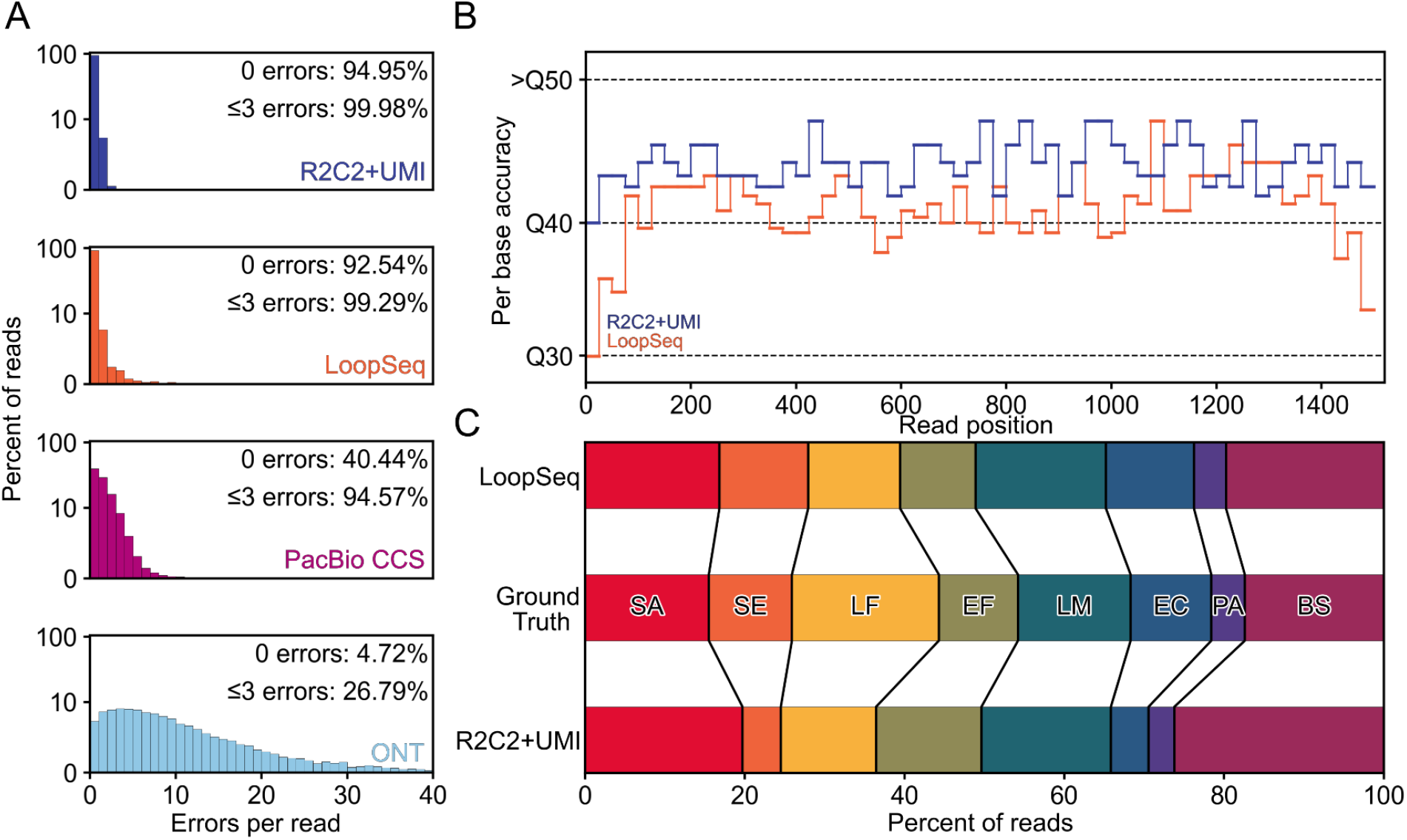
16S amplicon sequencing. A) Distribution of errors-per-read is shown for ONT raw reads, PacBio CCS reads, Loop-Seq reads, and R2C2+UMI reads, B) estimated Per base accuracy of R2C2+UMI and LoopSeq reads, and C) species composition covering 16S genes present in the ZymoBIOMICS Microbial Community DNA Standard are shown.

Third, to see whether the previously measured overall accuracies were constant throughout the amplicon, we measured position-dependent per-base accuracies of the two most accurate methods - R2C2+UMI and LoopSeq. LoopSeq accuracy remained between ∼Q39-Q44 throughout most of the amplicon but dipped to as low as Q30 in the first and last 50nt - possibly a result of their (proprietary) library preparation. In contrast, R2C2+UMI accuracy never dipped below Q40, instead remaining between ∼Q43-Q48 throughout most of the amplicon (Fig. 3B).

Overall, this suggested that using approximately 20% of a PromethION flow cell run-time, R2C2+UMI generated 2-times the reads at 3-times the accuracy of a publicly available LoopSeq dataset, all while capturing the overall diversity of the input sample (Fig. 3C).

## Discussion

When sequencing amplicons, achieving the highest possible sequencing accuracy removes experimental uncertainty from analysis and enables more meaningful interpretation of the resulting data. This applies to the IGH and 16S amplicons sequenced here but extends to other applications like the targeted sequencing of mixed cancer/normal samples to look for rare mutations. Illumina sequencing using UMIs has been the gold standard for amplicon sequencing^28^. However, Illumina sequencers are in fact ill-suited for amplicon sequencing, struggling with read length and library complexity. Despite this, there are a lot of protocols that produce Illumina-style amplicon libraries which would represent a significant resource if the limitations of Illumina sequencers could be overcome. We have previously shown that, using R2C2, <100ng of Illumina-style libraries can be converted and sequenced accurately and cost-effectively on ONT sequencers, immediately overcoming restrictive library complexity and length limitations^11^.

Here we showed that using R2C2 to convert Illumina-style, UMI-containing amplicon libraries and sequencing these converted libraries with R10.4 flowcells (R2C2+UMI) outperforms sequencing those same amplicon libraries on an Illumina MiSeq 2x300 (Illumina+UMI). At the same number of reads, R2C2+UMI achieved 100-fold higher consensus accuracy in the center of a ∼550nt IGH amplicon. Notably, R2C2+UMI also achieved 3-fold higher accuracy than synthetic long reads generated using the LoopSeq method when applied to 16S amplicons. In both cases, R2C2+UMI has set new benchmarks by achieving Q51.7 sequencing accuracies for IGH amplicons and ∼Q44.8 sequencing accuracies for 16S amplicons - to our knowledge the highest measured accuracies reported for these applications to date.

To achieve this level of accuracy, we developed and applied the easy-to-setup-and-use BC1 computational tool with two major considerations: Useability and scalability. To increase useability, BC1 was designed to require only a single command to find and parse UMIs, then to create very high accuracy consensus reads for each UMI. To enable scalability, BC1 was designed to require only a small amount of RAM, and to be multithreaded, with the slowest part of the pipeline - polishing with medaka - being optional.

This leaves cost as the final consideration when deciding to generate and analyze R2C2+UMI data instead of Illumina+UMI data. R2C2 reads, when generated using a PromethION flow cell, have comparable cost to Illumina MiSeq reads - the only Illumina sequencer capable of 2x300 paired end sequencing - a $900 PromethION flow cell generates between 8-15 million R2C2 reads, an $2500 2x300 MiSeq flow cell generates about 30 million paired end reads. Importantly, PromethION flow cells can now be processed on the P2Solo which carries very low capital cost, making it possible for most molecular biology labs to perform these experiments in their own labs. Additionally, because C3POa and BC1 workflows can be performed on consumer-grade computer workstations, analysis of the resulting data can be performed locally as well.

Overall, our results show that the R2C2+UMI approach is an ideal choice for sequencing amplicons, combining the highest possible accuracy with low cost. R2C2+UMI will therefore enable detailed biological insights by empowering ultra-accurate amplicon sequencing for a diverse array of researchers and research questions.

## Methods

### IGH

#### IGH amplicon generation

200ng of RNA extracted from whole human blood used as input for reverse transcription using Smartscribe RT (Takara Clontech) with primers targeting the IGH constant regions (IDT, Supplemental Table S1) (72°C for 3 minutes, 42°C for 60 minutes and 70°C for 5 minutes). A nested set of constant region specific primers and a set of primers V segment specific primers, both of which contain UMIs and partial p5 or p7 sequences were then used for a 2-cycle PCR using Phusion polymerase (ThermoFisher) to uniquely label each original IGH transcript (2 cycles of 98°C for 2 minutes, 52°C for 2 minutes, and 72°C for 4 minutes). The labeled molecules were then amplified using primers attaching complete p5 and p7 Illumina adapters to the molecules using KAPA HiFi HotStart ReadyMix (Roche) (95°C for 3 minutes followed by 25 cycles of 98°C for 20 seconds, 67°C for 15 seconds, and 72°C for 1 minutes, and last elongation at 72°C for 5 minutes).

#### IGH amplicon sequencing

UMI-labeled IGH amplicons were sequenced on multiplexed 2x300 flow cells on the Illumina MiSeq or converted to R2C2 libraries using the Illumina But With Nanopore workflow described previously^11^ and then sequenced on the ONT PromethION using LSK114 ligation kits and R10.4 flow cells.

#### IGH amplicon analysis

##### Illumina data

Paired end 2x300 Illumina reads were processed into full-length amplicon sequences using the pResto group of tools with the exception of UMI clustering, which was performed using our own IGH processing pipeline (https://github.com/christopher-vollmers/IGH_pipeline) to keep more consistent with the UMI clustering of the R2C2 data. For detailed commands, see supplemental information.

##### R2C2 data

ONT reads were basecalled using the R10.4 SUP model of guppy(v6). Raw reads were then processed and demultiplexed into R2C2 consensus reads and their subreads using C3POa. These R2C2 consensus reads and subreads were then processed into R2C2+UMI consensus reads using the following BC1 command:

~~~
python3 bc1.py -i IGH_C3POa.fasta -u 5.0:12.GACAG.NNNNNNNNNNNNNN.,3.0:12.GACAG.NNNNNNNNNNNNNN. –o Sample1.bd1 -s R2C2_subreads.fastq -f -t 60 –m
~~~

##### Error analysis

R2C2+UMI and Illumina+UMI reads were processed using IgBlast (v1.14). IgBlast assigned each read to V, D, and J segments (as downloaded from IMGT) and determined mismatches, and indels between the read and assigned segment. Separately, R2C2+UMI and Illumina+UMI reads were aligned to constant region exons using mappy to determine isotype and mismatches and indels in that region.

For error analysis within the constant regions counting mismatches/indels were simply counted for each read. For error analysis within V and J segments - to take somatic hypermutation into account, which is indistinguishable from sequencing error - R2C2+UMI and Illumina+UMI reads were matched into pairs based on their identical UMIs. For each pair, mismatch and indel positions as determined by IgBlast for R2C2+UMI and Illumina+UMI reads were compared between the reads. If one of the reads contained a mismatch or indel at a specific position and the other read did not, a sequencing error for the first technology was declared.

To adjust for read depth when visualizing position-dependent error rates, we calculated error rates for each method at each position and then simulated the number of errors likely to occur in exactly 100,000 bases at those error rates.

### 16S

#### 16S amplicon generation

100ng of ZymoBIOMICS Microbial Community DNA Standard (Zymo Research D6305) was used as template for 2-cycle PCR (Phusion polymerase - ThermoFisher) with 16S primers capturing the majority of the 16S gene and containing UMIs and partial p5 or p7 adapters (Supplemental Table S1). The resulting UMI-labeled 16S dsDNA was then amplified using indexed p5/p7 primers and KAPA HiFi HotStart ReadyMix (Roche) (95°C for 3 minutes followed by 25 cycles of 98°C for 20 seconds, 67°C for 15 seconds, and 72°C for 1 minutes, and last elongation at 72°C for 5 minutes).

#### 16S amplicon sequencing

UMI-labeled 16S amplicons were then converted to R2C2 libraries using the Illumina But With Nanopore workflow described previously with the modification that the splint was generated through the simple annealing of two oligos (Supplemental Table S1) instead of primer extension. and the sequencing using LSK114 ligation kits on R10.4 flow cells on the ONT P2Solo.

16S amplicon analysis

ONT reads were basecalled using the R10.4 SUP model of guppy(v6). Raw reads were then processed and into R2C2 consensus reads and their subreads using C3POa. These R2C2 consensus reads and subreads were then processed into R2C2+UMI consensus reads using the following BC1 command:

~~~
python3 bc1.py -i 16S_C3POa.fasta -o 16S.BC1_consensus -s R2C2_Subreads.fastq -f -t 60 -u 5.38:50.ACAG.BDHVBDHVBDHV.AG,3.38:50.ACAG.BDHVBDHVBDHV.CG
~~~

PacBio reads from^8^ were downloaded from the SRA at SRR9089357. LoopSeq reads from^5^ were downloaded from the SRA at SRR12148161.

Both PacBio and LoopSeq reads were filtered for perfect matches to the primers used in their respective studies and the primers were then trimmed.

##### Error analysis

Reads were aligned to the 16S genes extracted from the complete reference genomes of the bacteria present in the ZymoBIOMICS Microbial Community DNA Standard (Zymo Research D6305) using minimap2. Total errors were extracted from the resulting SAM file. Position dependent errors were determined by converting the resulting SAM file into pairwise alignments using sam2pairwise^29^ which were then parsed to catalog errors and their positions.

To adjust for read depth when visualizing position-dependent error rates, we calculated error rates for each method at each position and then simulated the number of errors likely to occur in exactly 100,000 bases at those error rates.

#### General

Samtools^30^, minimap2^31^, numpy^32^, scipy^33^, matplotlib^34^, sam2pairwise^29^ were used throughout the study.

## Code availability

The BC1 tool is available at github at https://github.com/christopher-vollmers/BC1. Tools for trimming sequencing primers and filtering chimeras used in the study can also be found in that repository

## Data availability

Fully processed and filtered IGH amplicon data is available from the Sequence Read Archive (SRA) at PRJNA1005293. Fully processed and filtered 16S amplicon data is available from the Sequence Read Archive (SRA) at PRJNA1005263. 16S sequence references for the ZymoBIOMICS Microbial Community DNA Standard (Zymo Research D6305) are provided as supplemental data file1.

## Acknowledgements

This work was supported by the NIH/NIGMS grant R35GM133569 to C.V.. This work was supported by NHLBI R01HL124233, U01 HL089897, R01 HL147326, and U01 HL089856. The COPDGene study (NCT00608764) is also supported by the COPD Foundation through contributions made to an Industry Advisory Board that has included AstraZeneca, Bayer Pharmaceuticals, Boehringer-Ingelheim, Genentech, GlaxoSmithKline, Novartis, Pfizer, and Sunovion.

## Supplemental Information

### Supplemental methods

Presto commands for the parsing and consensus generation of MiSeq 2x300 IGH Illumina data starting with demultiplexed Sample1_R1.fastq and Sample_R2.fastq paired reads.

~~~
FilterSeq.py quality -s Sample1_R1.fastq -q 10 --outname R1 --log FS1.log
FilterSeq.py quality -s Sample1_R2.fastq -q 10 --outname R2 --log FS2.log
~~~

~~~
ParseLog.py -l FS1.log FS2.log -f ID QUALITY
~~~

~~~
MaskPrimers.py score -s R1_quality-pass.fastq -p IGHV_Primers.fasta --start 14 --mode cut –barcode --outname R1 --log MP1.log
MaskPrimers.py score -s R2_quality-pass.fastq -p IGHC_Primers.fasta --start 14 --mode cut –barcode --outname R2 --log MP2.log
~~~

~~~
ParseLog.py -l MP1.log MP2.log -f ID PRIMER BARCODE ERROR
~~~

~~~
PairSeq.py -1 R1_primers-pass.fastq -2 R2_primers-pass.fastq --1f BARCODE --2f BARCODE --coord presto --act cat
~~~

### Custom_script to add UMI_indexes

~~~
python3 add_UMI_index.py indexed_reads_mismatch_sorted.txt R1_primers-pass_pair-pass.fastq R1_primers-pass_pair-pass_indexed.fastq
python3 add_UMI_index.py indexed_reads_mismatch_sorted.txt R2_primers-pass_pair-pass.fastq R2_primers-pass_pair-pass_indexed.fastq
~~~

~~~
AlignSets.py muscle -s R1_primers-pass_pair-pass_indexed.fastq --bf UMI_INDEX --exec muscle --outname R1 --log AS1.log
AlignSets.py muscle -s R2_primers-pass_pair-pass_indexed.fastq --bf UMI_INDEX --exec muscle --outname R2 --log AS2.log
~~~

~~~
BuildConsensus.py -s R1_aligned-pass.fastq --bf UMI_INDEX --pf PRIMER --prcons 0.6 --maxerror 0.1 –maxgap 0.5 --outname R1 --log BC1.log
BuildConsensus.py -s R2_aligned-pass.fastq --bf UMI_INDEX --pf PRIMER --prcons 0.6 --maxerror 0.1 –maxgap 0.5 --outname R2 --log BC2.log
~~~

~~~
ParseLog.py -l BC1.log BC2.log -f BARCODE SEQCOUNT CONSCOUNT PRIMER PRCONS PRCOUNT UMI_INDEX
~~~

~~~
PairSeq.py -1 R1_consensus-pass.fastq -2 R2_consensus-pass.fastq --coord presto
~~~

~~~
AssemblePairs.py align -1 R2_consensus-pass_pair-pass.fastq -2 R1_consensus-pass_pair-pass.fastq --coord presto --rc tail --1f CONSCOUNT --2f CONSCOUNT PRCONS --outname Sample1 --log AP.log
~~~

This results in a `Sample1_assemble-pass.fastq` file which is processed further in downstream analysis.

**Table S1:**
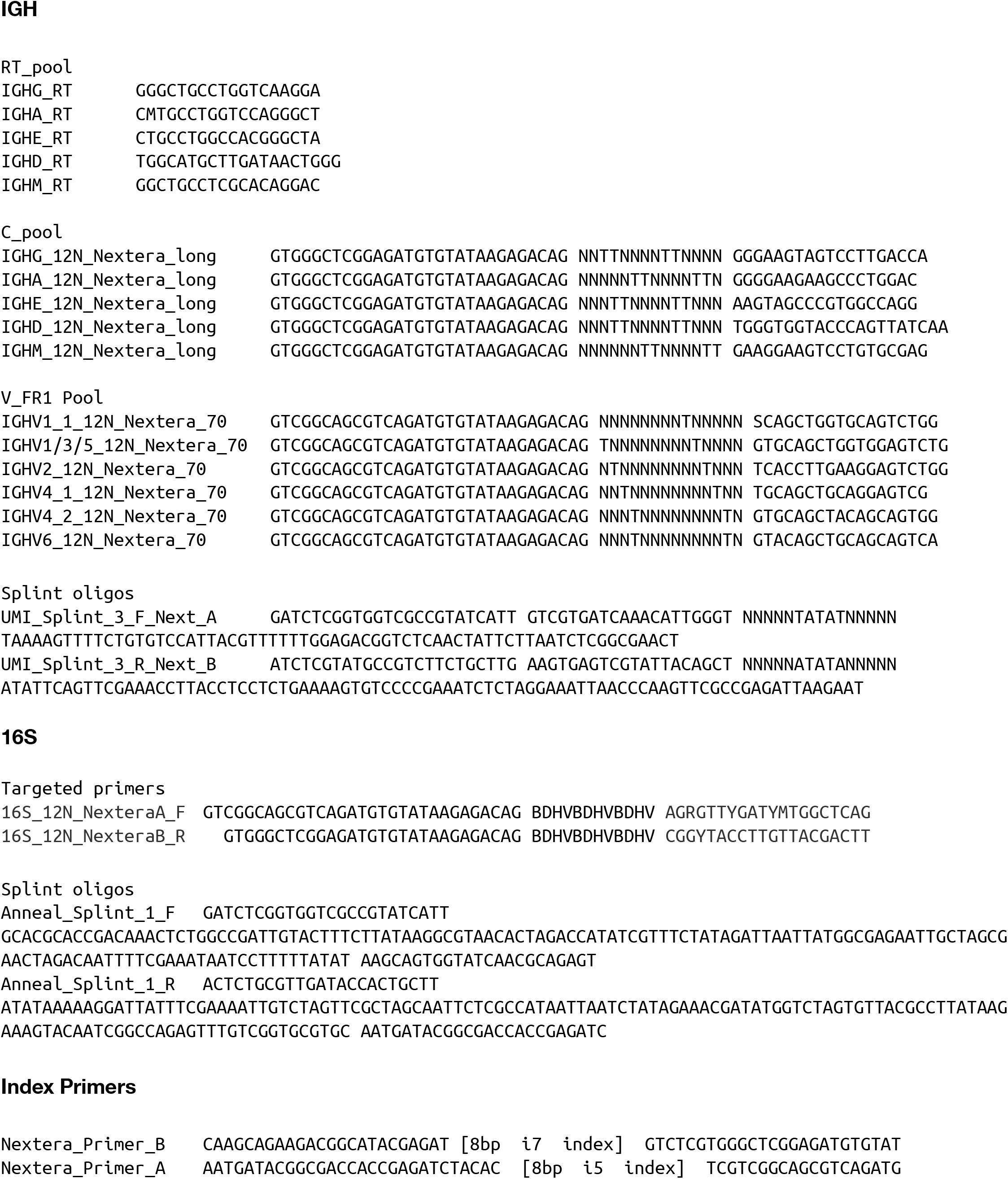
Oligos used in this study

